# Sexual dimorphism in epilepsy and comorbidities in Dravet syndrome mice carrying a targeted deletion of exon 1 of the *Scn1a* gene

**DOI:** 10.1101/2021.08.27.457904

**Authors:** Rogério R. Gerbatin, Joana Augusto, Halima Boutouil, Cristina R. Reschke, David C. Henshall

**Author notes:** **Corresponding author:** Department of Physiology & Medical Physics, RCSI University of Medicine and Health Sciences, 123 St. Stephen’s Green, Dublin D02 YN77, Ireland. E-mail address (D.C. Henshall).

## Abstract

**Objective:** Dravet Syndrome (DS) is a catastrophic form of paediatric epilepsy associated with multiple comorbidities mainly caused by mutations in the *SCN1A* gene. DS progresses in three different phases termed febrile, worsening and stabilization stage. Mice that are haploinsufficient for *Scn1a* faithfully model each stage of DS, although various aspects have not been fully described, including the temporal appearance and sex differences of the epilepsy and comorbidities. The aim of the present study was to investigate the epilepsy landscape according to the progression of DS and the long-term co-morbidities in the *Scn1a(+/-)*^tm1Kea^ DS mouse line that are not fully understood yet.

**Methods:** Male and female F1.*Scn1a(+/+)* and F1.*Scn1a(+/-)*^tm1Kea^ mice were assessed in the hyperthermia model or monitored by video electroencephalogram (vEEG) and wireless video-EEG according to the respective stage of DS. Long-term comorbidities were investigated through a battery of behaviour assessments in ∼6 month-old mice.

**Results:** At P18, F1.*Scn1a(+/-)*^tm1Kea^ mice showed the expected sensitivity to hyperthermia-induced seizures. Between P21 and P28, EEG recordings in F1.*Scn1a(+/-)*^tm1Kea^ mice combined with video monitoring revealed a high frequency of SRS and SUDEP. Power spectral analyses of background EEG activity also revealed that low EEG power in multiple frequency bands was associated with SUDEP risk in F1.*Scn1a(+/-)*^tm1Kea^ mice during the worsening stage of DS. Later, SRS and SUDEP rates stabilized and then declined in F1.*Scn1a(+/-)*^*tm1kea*^ mice. SRS and SUDEP in F1.*Scn1a(+/-)*^*tm1kea*^ mice displayed variations with the time of day and sex, with female mice displaying higher numbers of seizures and greater SUDEP risk. F1.*Scn1a(+/-)*^*tm1kea*^ mice ∼6 month-old displayed fewer behavioural impairments than expected including hyperactivity, impaired exploratory behaviour and poor nest building performance.

**Significance:** These results reveal new features of this model that will optimize use and selection of phenotype assays for future studies on the mechanisms, diagnosis, and treatment of DS.

**Key point box:**

- *Scn1a(+/-)*^*tm1kea*^ DS mouse model faithfully reproduces the three stages of DS
- Sex of F1.*Scn1a(+/-)*^*tm1kea*^ mice influences the epilepsy phenotype
- F1.*Scn1a(+/-)*^*tm1kea*^ develop some of the long-term comorbidities of DS

## 1 INTRODUCTION

Dravet Syndrome (DS) is a rare, intractable and catastrophic form of childhood epilepsy with an estimated incidence of 1 in 17,700 to 40,000 births.^1, 2^ Nearly 90% of DS patients carry *de novo* heterozygous mutations in the *SCN1A* gene leading to haploinsufficiency of the type 1 voltage-gated sodium channel α subunit (Nav1.1).^3^ Loss of Nav1.1 protein mainly affects parvalbumin-expressing interneurons, causing disruption of excitation and inhibition balance in several neuronal circuits.^4^ Recent studies show this impairment is transient, however, and the mechanisms by which *SCN1A* deficiency contributes to cognitive and other phenotypes remains incompletely understood.^5, 6^

DS is primarily characterized by hyperthermia sensitivity, spontaneous recurrent seizures (SRS) and premature death. Apart from epilepsy, a global development delay, hyperactivity, intellectual disability, and autistic like-behaviour may be also present.^7^ The behavioural impairments and epilepsy phenotype emerge in an age-dependent manner according to three different stages of DS termed ‘febrile’, ‘worsening’ and ‘stabilization’ stage.^8^ Thus, a comprehensive understanding of DS features in each disease stage is crucial for successful preclinical development and evaluation of new targeted therapies or biomarker discovery.

Several genetic mouse lines have been developed to model DS.^4, 9, 10^ This includes the *Scn1a*^tm1Kea^ mouse line harbouring a targeted deletion of exon 1 of *Scn1a*.^11^ This model has demonstrated translational value in identifying known and novel anti-seizure molecules and has helped define the molecular landscape upon Scn1a deletion.^12^ While the broad features of the model are resolved, various phenotypes remain incompletely defined. This includes epilepsy-related features of *Scn1a(+/-)*^tm1Kea^ mice at the different stages of DS progression and the influence, if any, of sex. Furthermore, to the best of our knowledge, long-term assessment of neuropsychiatric comorbidities of this Dravet mouse line have not been evaluated.

Here we sought to characterise the epilepsy-related features according to the progression of DS in F1.*Scn1a(+/-)*^tm1Kea^ mice. Moreover, we investigated the long-term behaviour impairments to characterize the comorbidities of the F1.*Scn1a(+/-)*^tm1Kea^ DS mouse line. Our findings uncover previously unknown features. We identify two seizure types in F1.*Scn1a(+/-)*^tm1Kea^ subject to hyperthermia at P18. Sex differences and the time of the day may influence the occurrence of SRS and SUDEP in F1.*Scn1a(+/-)*^tm1Kea^ mice. Finally, long-term comorbidities in F1.*Scn1a(+/-)*^tm1Kea^ mice are characterized by hyperactivity, impaired exploratory behaviour, poor nesting building ability but not changes in memory performance, sociability or anxiety levels.

## 2. MATERIALS AND METHODS

### 2.1 Mice and ethics statement

All animal experiments were performed in accordance with the European Communities Council Directive (86/609/EEC) and approved by the Research Ethics Committee of the Royal College of Surgeons in Ireland (REC 1302bbb) under license from the Department of Health (AE19127/P064), Dublin Ireland. Animals were maintained in a light (08:00 - 20:00) / dark cycle (20:00 - 08:00) with food and water *ad libitum*. F1.*Scn1a(+/-)*^tm1Kea^ mice used in the experiments were generated by crossing *Scn1a(+/-)*^tm1Kea^ male on the 129S6/SvEvTac background (Jackson Laboratory, USA) with inbred female mice C57BL/6JOlaHsd (Envigo, UK).^11^

### 2.2 Experimental design

All animals were genotyped before P7 and then assigned to four different time points to investigate the three different stages of DS: Stage 1 (Febrile): At P18, the first cohort of mice were subjected to a hyperthermia-induced seizure threshold assay to determine their sensitivity to febrile seizures. Stage 2 (Worsening): At P21, a second cohort of mice were implanted with cortical EEG electrodes to be recorded by vEEG followed by video monitoring until P28 to investigate SRS and SUDEP. Stage 3 (Stabilization): At P36, another group of mice were implanted with a wireless telemetry device for continuous vEEG recording to detect SRS and SUDEP for two weeks until P49. Finally, another cohort of naïve mice were monitored from P0 to 6 months of age for the occurrence of SUDEP. Animals that survived and completing 5 – 6 months old were subject to a battery of behavioural assessments to investigate the long-term comorbidities in DS (Figure 1).

**FIGURE 1.**
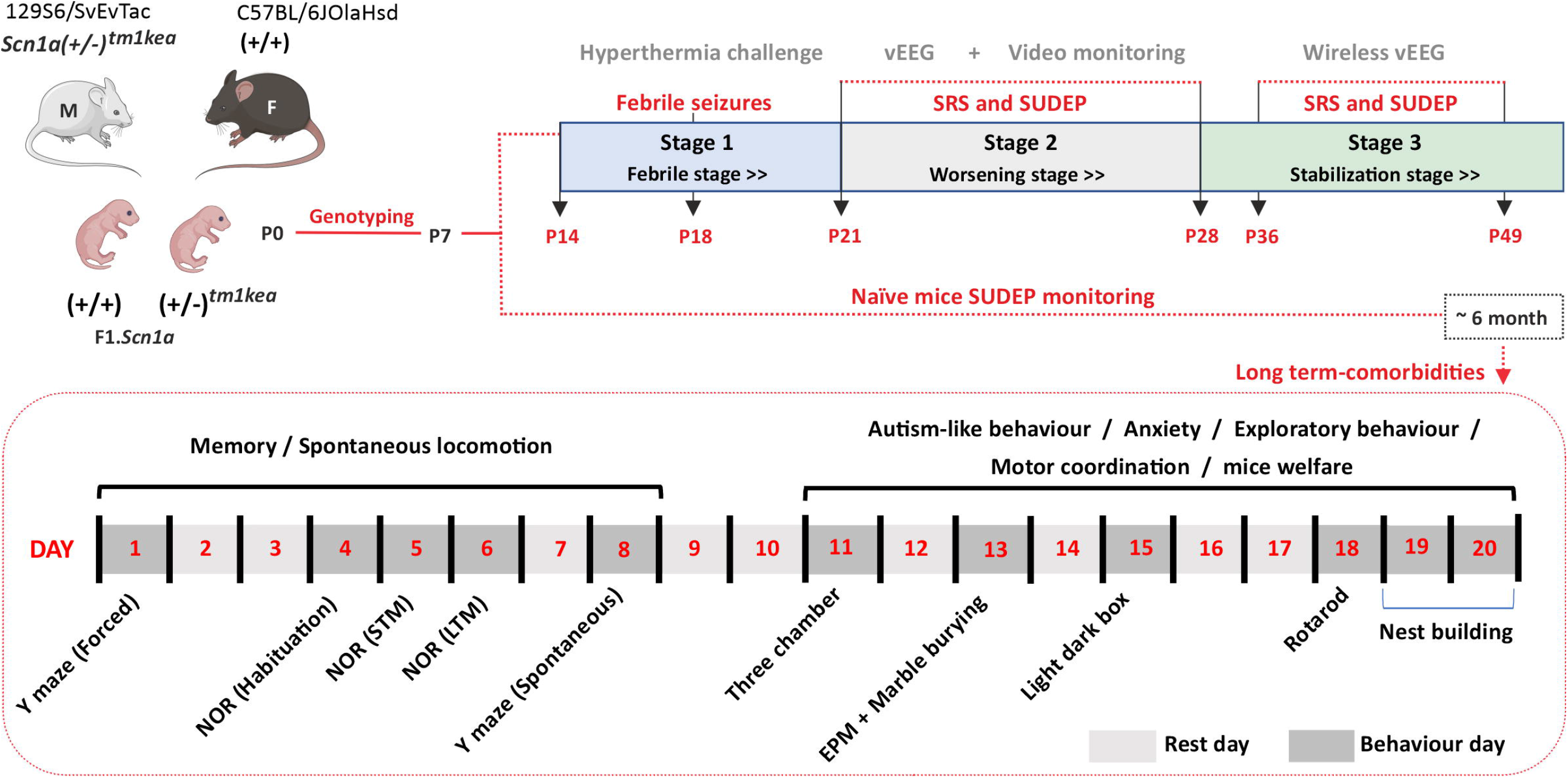
Experimental design. Overview of the different experiments and analyses. Figure drawn using Biorender.

### 2.4 Seizure semiology

All seizures detected in each stage of DS were scored according to the Racine scale with a few modifications, as reported:^13, 14^ No behaviour changes (0), mouth and facial movements (1), head nodding (2), unilateral forelimb clonus (3), bilateral forelimb clonus with rearing (4), rearing and falling (loss of posture) (5), wild running or jumping (6) and Tonic hindlimb extension possibly leading to death (7).

### 2.4 Behaviour phenotype

Naïve 5 to 6-month-old mice were submitted to a battery of behavioural assessments. Spatial reference, working and recognition memory was assessed with the Y maze and novel object recognition test, respectively^15, 16^ Spontaneous locomotion, stereotyped behaviour, and anxiety-like behaviour was investigated in the open field. Further investigation related to anxiety-like behaviour was performed in the elevated plus maze and light dark box test as previously described^17^. Autism-like behaviour and motor coordination were assessed in the three chamber and rotarod test, respectively.^18, 19^ Finally, innate exploratory behaviour and welfare parameters were assessed through marble burying and the nest building task.^20, 21^ Behavioural tests were performed with multiple rest day intervals and according to the increasing order of interventional complexity.

### 2.3 Statistical analyses

The normality of the data was analysed using D’Agostino and Pearson’s omnibus normality test. Data were analysed using unpaired two-tailed Student t test, Mann-Whitney U test, Kaplan-Meier method, Spearman’s rank-order correlation, Two-way repeated measures ANOVA, one and two-way analysis of variance (ANOVA) or permutation test followed by Tukey’s post hoc test, as appropriate. The specific statistical test used for each experiment are indicated in the figure legends. Data are expressed as mean ± SEM or median with interquartile range, as appropriate. Differences between groups were considered statistically significant when p<0.05. Experiments and data were analysed blind to genotype.

## 3. RESULTS

### 3.1 F1.*Scn1a(+/-)*^*tm1kea*^ P18 mice display sensitivity to hyperthermia-induced seizures during the febrile stage of DS

Febrile seizures as a result of sensitivity to hyperthermia are a hallmark of DS onset. Thus, we first investigated whether F1.*Scn1a(+/-)*^*tm1kea*^ mice display sensitivity to hyperthermia-induced seizures at an early age (P18). Figure 2 shows the susceptibility of F1.*Scn1a(+/-)*^*tm1kea*^ mice to hyperthermia-induced seizures. As body temperature was elevated, all F1.*Scn1a(+/-)*^*tm1kea*^ animals manifested clinically recognizable seizures at temperatures ranging from 39.4°C to 41.2°C (Figure 2A). In contrast, F1.*Scn1a(+/+)* mice did not experience seizures at any body temperature up to the cut-off (maximum) temperature of 42.5°C. Interestingly, we noted seizures fell into two distributions based on duration in F1.*Scn1a(+/-)*^*tm1kea*^ mice. Half (50%) of F1.*Scn1a(+/-)*^*tm1kea*^ exhibited seizures lasting 10 to 14s whereas the remaining 50% displayed longer seizures around 20s (Figure 2B) (Movies S1,S2). The seizure duration sub-phenotypes did not correlate with the threshold temperature for seizures during the hyperthermia challenge (Figure S1). Most seizures in F1.*Scn1a(+/-)*^*tm1kea*^ mice reached severity 5 without progressing to wild jumping/running, hindlimb extension (Figure 2C) or sudden death (Figure 2D). Analysis of male and female F1.*Scn1a(+/-)*^*tm1kea*^ mice revealed there were no significant sex differences in any of these parameters.

**FIGURE 2.**
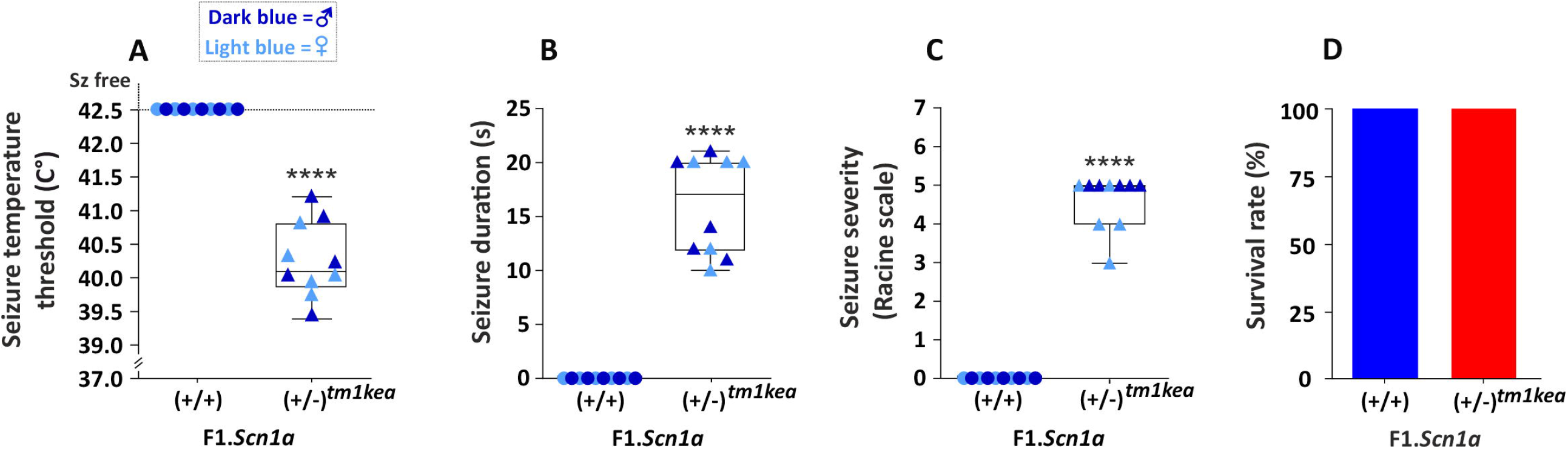
Characteristics of hyperthermia-induced seizures in F1.*Scn1a(+/-)*^*tm1kea*^ mice the during febrile stage of DS. Hyperthermia-induced seizures were induced in P18 male and female F1.*Scn1a(+/+)* or F1.*Scn1a(+/-)*^*tm1kea*^ mice (n = 10/group) by placing them in a thermostat-controlled heating chamber. A, as the temperature was raised, all F1.*Scn1a(+/-)*^*tm1kea*^ developed tonic-clonic seizures, with an onset averaging ∼40°C whereas no F1.*Scn1a(+/+)* mice developed seizures up to the cut-off temperature of 42.5°C. B, seizures in F1.*Scn1a(+/-)*^*tm1kea*^ mice lasted ∼17 seconds and C, averaged a score of 3-5 on a 7-point mouse-adapted Racine scale. D, no deaths occurred during or following seizures. No differences were observed between sexes (A-D). Mann-Whitney test ****p<0.0001.

### 3.2 F1.*Scn1a(+/-)*^*tm1kea*^ mice develop frequent and severe SRS during the worsening stage of DS

Within weeks of febrile seizure symptoms, Dravet patients experience SRS with increasing severity and frequency, reflecting the worsening stage (stage 2) of DS. Figure 3 shows the frequency, duration and severity of SRS observed during vEEG recordings and video-only monitoring from P21 to P28. F1.*Scn1a(+/-)*^*tm1kea*^ mice showed an increasing SRS frequency ranging from 4 to 20 seizures, often accompanied by death (SUDEP), whereas no seizures were observed in F1.*Scn1a(+/+)* control mice (Figure 3A,B). F1.*Scn1a(+/-)*^*tm1kea*^ mice displayed SRS with a median duration of 32s without a statistical difference between sexes (Figure 3C). However, analysis of the number of SRS in F1.*Scn1a(+/-)*^*tm1kea*^ mice revealed a significant sex difference, with more seizures in females (Figure 3D). Seizure severity in F1.*Scn1a(+/-)*^*tm1kea*^ mice showed an almost equal proportion of seizures scoring severity 5,6 and 7 on the adapted Racine Scale (Figure 3E). Spontaneous seizures were least severe at P21. From P22 to P25 F1.*Scn1a(+/-)*^*tm1kea*^ mice experienced the most severe SRS (Figure 3F). A sex difference was again observed. The total number of SRS with severity 6 and 7 observed in females F1.*Scn1a(+/-)*^*tm1kea*^ mice was twice that in males (Figure 3G). We also observed that severity 6 and 7 SRS were of a longer duration than SRS with severity 5 (Figure 3H), and a strong positive correlation was found between SRS duration and seizure severity (Figure 3I).

**FIGURE 3.**
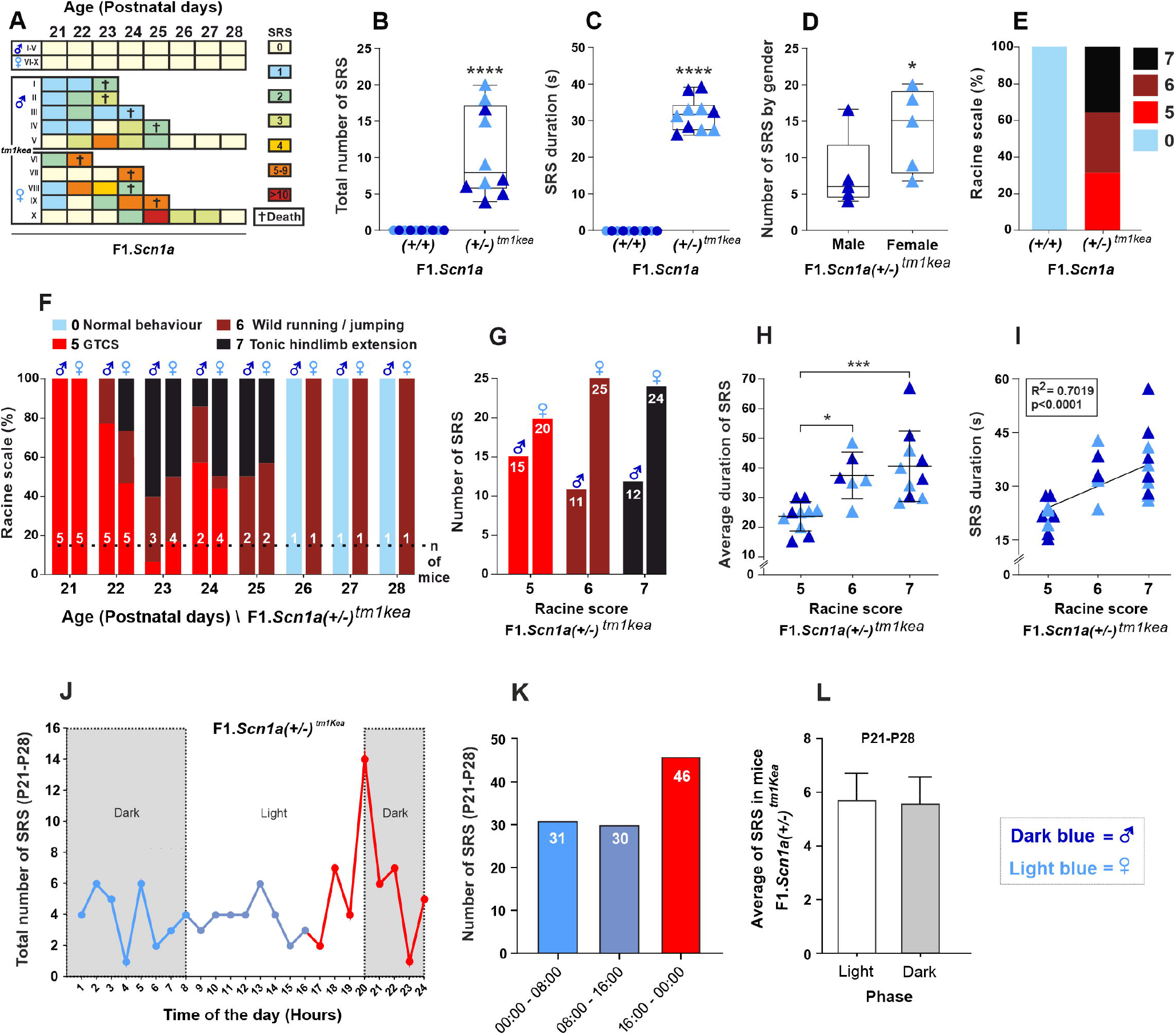
Features of SRS during the worsening stage of DS in F1.*Scn1a(+/-)*^*tm1kea*^ mice. A,B, Surface EEG combined with continuous video monitoring of male and female F1.*Scn1a(+/+)* or F1.*Scn1a(+/-)*^*tm1kea*^ mice (n = 10/group) between P21 – P28 detected the occurrence of SRS in all mice whereas no SRS were observed in F1.*Scn1a(+/+)* mice. EEG-confirmed SRS were always accompanied by clinical seizures. C, SRS in F1.*Scn1a(+/-)*^*tm1kea*^ lasted ∼ 30 seconds and D, female F1.*Scn1a(+/-)*^*tm1kea*^ mice experienced a higher number of SRS than male F1.*Scn1a(+/-)*^*tm1kea*^ mice. E, in general, F1.*Scn1a(+/-)*^*tm1kea*^ mice presented almost equal proportion of seizures scoring 5,6 and 7- point mouse-adapted Racine scale in which F, the most severe SRS were observed between P22 to P25. G, the number of SRS with severity 6 and 7 according to the adapted Racine scale was twice as high in female F1.*Scn1a(+/-)*^*tm1kea*^ mice when compared to male. F, SRS with severity 6 and 7 on the adapted Racine scale respectively showed a longer duration than seizures with severity 5 indicating I, a strong positive correlation between seizure duration and severity. J, F1.*Scn1a(+/-)*^*tm1kea*^ mice experience a peak of SRS between 19:00 −20:00. K, Number of SRS according to 8 hrs segment of the day was higher between 16:00 to 00:00 with L, no influence of light or dark phase in the incidence of SRS in F1.*Scn1a(+/-)*^*tm1kea*^ mice. *p<0.05, ***p<0.001, ****p<0.0001. B,C (Mann-Whitney test), D (Permutation test), H (one-way ANOVA), I (Spearman rank-order correlation), L (Student’s t-test).

Next, we investigated the time of day during which SRS were most likely to occur. Figure 3J shows a plot of 107 SRS recorded in F1.*Scn1a (+/-)*^*tm1kea*^ mice during monitoring between P21 and P28 according to the time of day. Interestingly, the number of SRS peaked just before the light-dark cycle lights went off (20:00 h). Dividing the day into 8 h segments also revealed that the number of SRS between 16:00-00:00 were 1.48 fold higher than those between 00:00-08:00 and 1.53 fold higher than between 00:08-16:00 (Figure 3K). Despite these patterns, the average number of SRS over the course of a full day did not differ between light and dark phases (Figure 3L).

### 3.3 F1.*Scn1a(+/-)*^*tm1kea*^ mice display higher incidence of SUDEP and reduction in background EEG power during the worsening stage of DS

SUDEP is a prominent feature of DS especially during the worsening stage of the disease (stage 2). A Kaplan-Meier plot of deaths revealed that F1.*Scn1a (+/-)*^*tm1kea*^ mice display a critical period of SUDEP risk from P22 to P25, with a survival rate of 20% at P28 (Figure 4A). There was no sex difference in the occurrence of SUDEP in F1.*Scn1a(+/-)*^*tm1kea*^ EEG-implanted mice between P21 to P28 (Figure 4B). All sudden deaths experienced by F1.*Scn1a(+/-)*^*tm1kea*^ mice were preceded by a severe SRS following a stereotypical progression. All pre-SUDEP SRS began with forelimb clonus, rearing and loss of balance/posture (GTCS-Racine scale 5) followed by wild running or jumping (Racine scale 6) ending with tonic hindlimb extension and possibly death (Racine scale 7) (Movie S3). Interestingly, the number of hindlimb extension seizures between 16:00-00:00 were 2.25 fold higher than those between 08:00-16:00 and 1.80 fold higher than between 00:00-08:00 (Figure 4C). Consistently, SUDEP incidence was also higher in the same period of day (16:00-00:00) being 2.5 fold higher than those between 08:00-16:00 and 5 fold higher than between 00:00-08:00 (Figure 4D). EEG power spectral analyses in the interictal period revealed low EEG power in multiple frequency bands in the period of high SUDEP incidence from P22 to P24/P25 in F1.*Scn1a(+/-)*^*tm1kea*^ mice (Figure 4E,F,G). Consistently, no SUDEP or reduction in EEG background power was observed at P21 in F1.*Scn1a(+/-)*^*tm1kea*^ mice. Figure 4 H,I shows representative EEG traces at P22 of F1.*Scn1a(+/+)* and F1.*Scn1a(+/-)*^*tm1kea*^ mice respectively. As can been observed, SUDEP in F1.*Scn1a(+/-)*^*tm1kea*^ mice was preceded by a severe SRS (ending with hindlimb extension).

**FIGURE 4.**
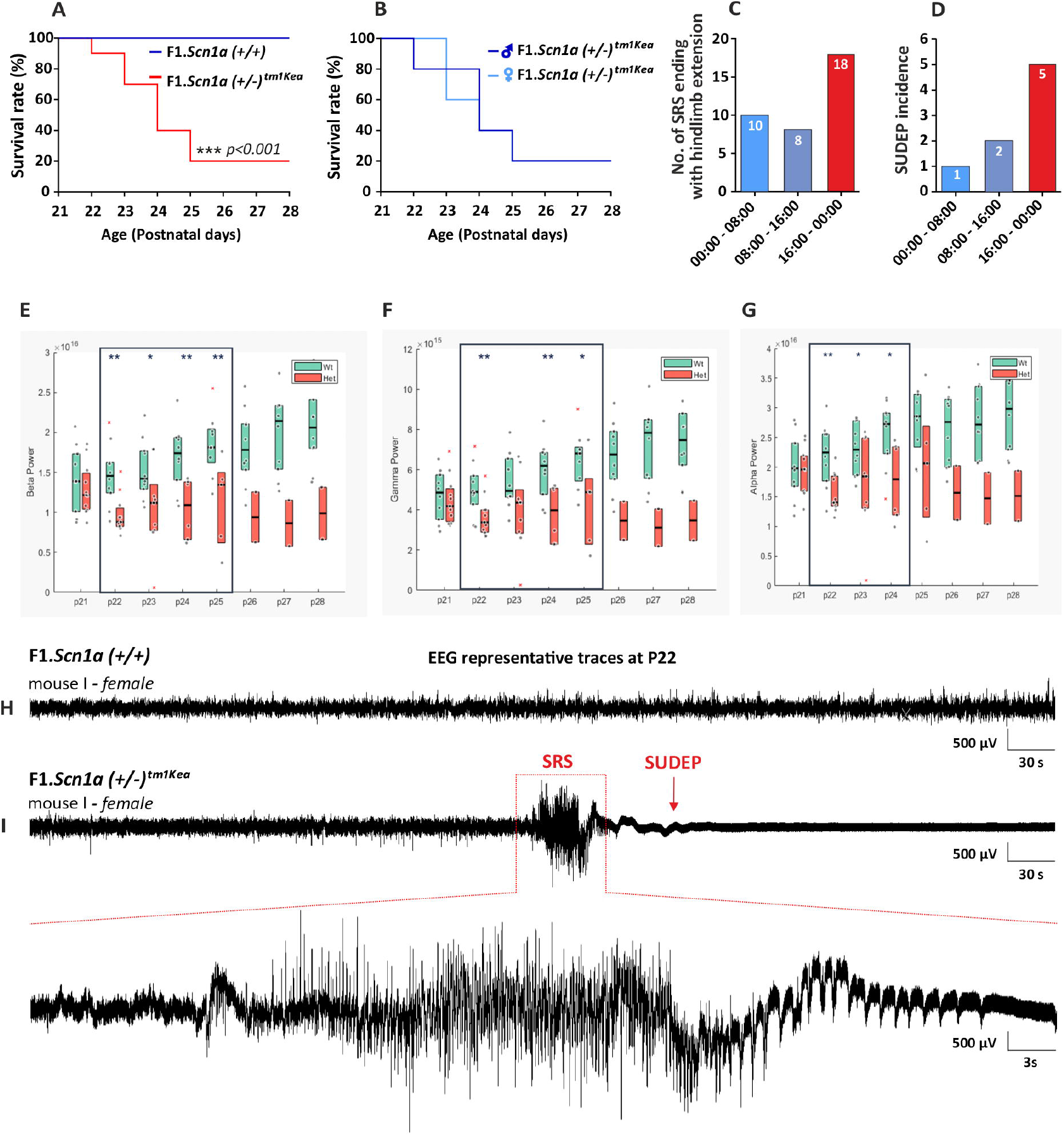
SUDEP during the worsening stage of DS in F1.*Scn1a(+/-)*^*tm1kea*^ mice. A, Surface EEG combined with continuous video monitoring of male and female F1.*Scn1a(+/+)* or F1.*Scn1a(+/-)*^*tm1kea*^ mice (n = 10/group) between P21 – P28 detected a SUDEP rate of 80% in F1.*Scn1a(+/-)*^*tm1kea*^ mice with B, no difference between male or female mice. C, Number of hindlimb extension seizures in F1.*Scn1a(+/-)*^*tm1kea*^ mice was more frequent between 16:00-00:00, in which D, SUDEP incidence was also higher in the same period of the day. E,F,G, F1.*Scn1a(+/-)*^*tm1kea*^ mice present low EEG power in multiple frequency bands when compared to F1.*Scn1a(+/+)* mice. I, EEG representative trace of F1.*Scn1a(+/+)* mice at P22 showing no SRS. J, EEG representative trace of F1.*Scn1a(+/-)*^*tm1kea*^ mice at P22 showing that SUDEP was preceded by a severe SRS. *p<0.05, **p<0.01, ***p<0.001. A,B (Log-rank Mantel-Cox test), E,F,G (Permutation test).

### 3.4 Reduced SRS and SUDEP incidence during the stabilization stage in F1.*Scn1a(+/-)*^tm1kea^ mice

SRS and SUDEP tend to reduce across adulthood during the stabilization stage of DS.^8, 22, 23^ Thus, we next equipped F1.*Scn1a* mice with implantable EEG telemetry units to track the occurrence of SRS and SUDEP in young adult mice (P36-P49). Monitoring during this period detected SRS in only 1 out of 5 F1.*Scn1a(+/-)*^*tm1kea*^ mice (Figure 5A). As expected, no seizures were observed in F1.*Scn1a(+/+)* control mice. Furthermore, all SRS observed over this period were less severe and did not reach the maximum score on the Racine scale (Figure 5B) and no deaths were recorded (Figure 5C). Figure 5D,E and F show representative EEG traces of one F1.*Scn1a(+/+)*, one F1.*Scn1a(+/-)*^*tm1kea*^ mice (without SRS) and one F1.*Scn1a(+/-)*^*tm1kea*^ mouse experiencing a SRS respectively. Later, using another batch of animals, we investigated the long-term survival in naïve mice. A Kaplan Meier plot revealed that the premature death commences at P20 culminating in overall mortality of ∼40% at the end of 6 months (Figure 5G). More than half of the deaths (56.25%) occurred within a short interval between P20 to P32 (worsening stage). Thereafter, the occurrence dropped, tending to stabilize over the period until 6 months (Figure 5G). Interestingly, a sub-group analysis of SUDEPs found a sex difference with higher mortality rates in female F1.*Scn1a(+/-)*^*tm1kea*^ mice (Figure 5G).

**FIGURE 5.**
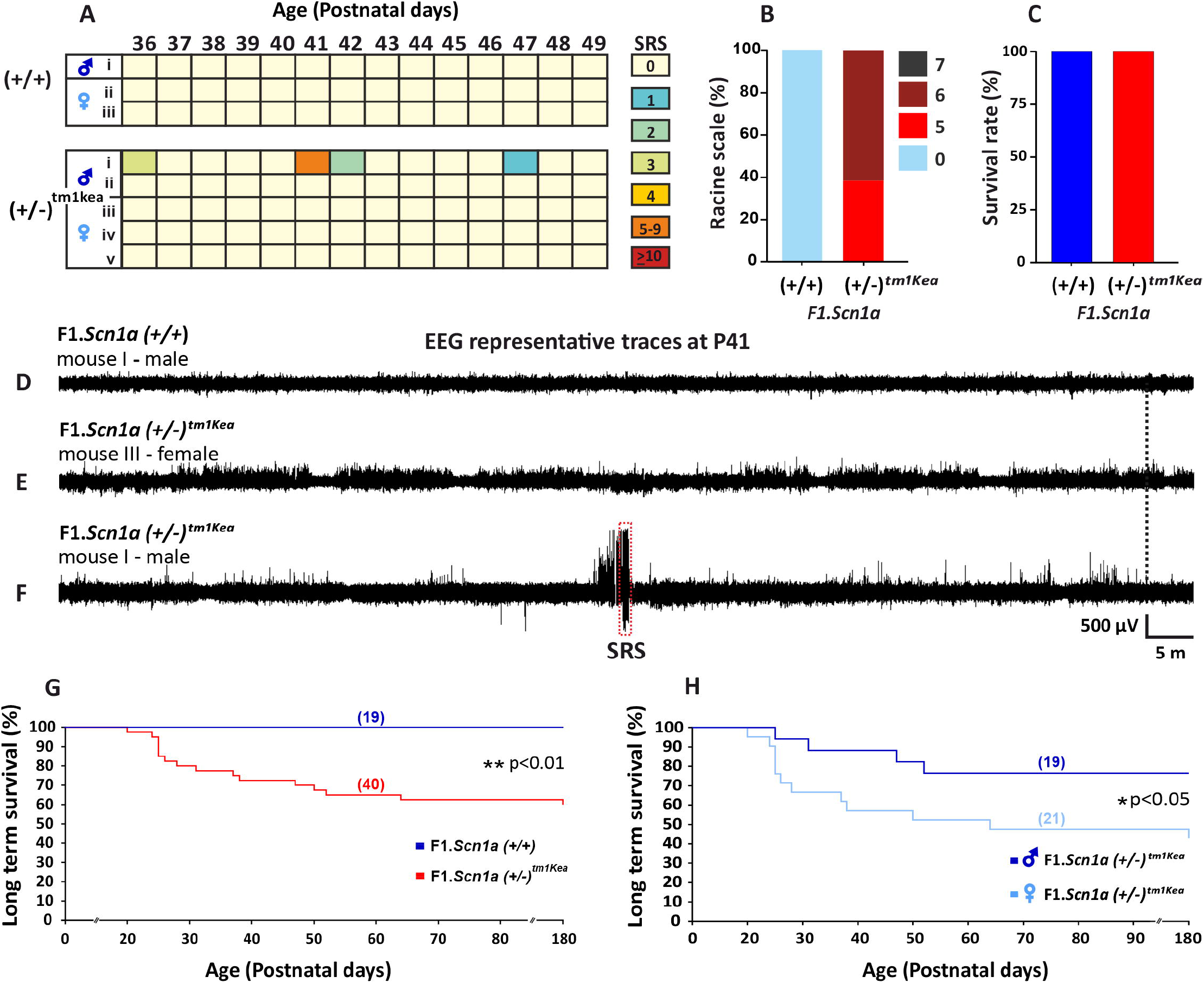
Characterization of SRS and SUDEP during the stabilization stage of DS in *F1*.*Scn1a(+/-)*^*tm1kea*^ mice. A,B, Video EEG monitoring confirmed that SRS are less frequent and severe as well C, SUDEP were rare in F1.*Scn1a(+/-)*^*tm1kea*^ mice (n=5) while no SRS were observed in F1.*Scn1a(+/+)* mice (n=3) (P36-P49). D, EEG representative trace of F1.*Scn1a(+/+)* mice and E, F1.*Scn1a(+/-)*^*tm1kea*^ mice showing no EEG seizure. F, EEG representative trace of F1.*Scn1a(+/-)*^*tm1kea*^ mice displaying a SRS. G, Monitoring of naïve male and female F1.*Scn1a(+/+)*(n=19) or F1.*Scn1a(+/-)*^*tm1kea*^ mice (n = 38) between P0 – 6 month old confirmed that more than 50% of F1.*Scn1a(+/-)*^*tm1kea*^ mice experience SUDEP from between P20 to P32 which later on tend to stabilize. H, female F1.*Scn1a(+/-)*^*tm1kea*^ mice displayed higher number of SUDEP when compared to male F1.*Scn1a(+/-)*^*tm1kea*^ mice. Log-rank Mantel-Cox test *p<0.05, **p<0.01.

### 3.5 Long-term neuropsychiatric comorbidities in F1.*Scn1a(+/-)*^*tm1kea*^ mice

Figure 6 shows the long-term comorbidities in ∼6 month old F1.*Scn1a(+/-)*^*tm1kea*^ mice. F1.*Scn1a(+/-)*^*tm1kea*^ mice displayed increased spontaneous locomotion activity in the open field test, displaying an increase in total distance travelled (Figure 6A), acceleration (Figure 6B), speed (Figure 6C) and total time mobile (Figure 6D), when compared to F1.*Scn1a(+/+)* control mice (Figure 6E). Assays of autism-like behaviour revealed that F1.*Scn1a(+/-)*^*tm1kea*^ mice display more stereotyped behaviour (repetitive rotations of animal’s body) in the open field test (Figure 6F). F1.*Scn1a(+/-)*^*tm1kea*^ mice displayed normal social interaction (Figure 6G,I) and response to social novelty (Figure 6H,I) in the three chamber test. Similarly, F1.*Scn1a(+/-)*^*tm1kea*^ mice did not show anxiety-like behaviour in peripheral regions of the open field test (Figure S2), elevated plus maze (Figure 6J) or light dark box (Figure 6K), when compared to F1.*Scn1a(+/+)* mice. Interestingly, F1.*Scn1a(+/-)*^*tm1kea*^ mice also performed normally when compared to F1.*Scn1a(+/+)* mice in three memory paradigms: Spatial reference memory (y maze forced, Figure 6L), spatial working memory (y maze spontaneous, Figure 6M) and recognition memory (novel object recognition, Figure 6N). Finally, we investigated the exploratory behaviour and nest building activity in the marble burying and nest build task, respectively. F1.*Scn1a(+/-)*^*tm1kea*^ mice showed a reduction in the number of marbles buried (Figure 6P,R) and displaced (Figure 6Q,R), as well as poor nest-building ability at 24 h (Figure 6O) and 48 h (Figure 6O) when compared to F1.*Scn1a(+/+)* mice). Analysis of male and female F1.*Scn1a(+/-)*^*tm1kea*^ mice revealed no significant difference in any behaviour parameters (Figure S2-S3).

**FIGURE 6.**
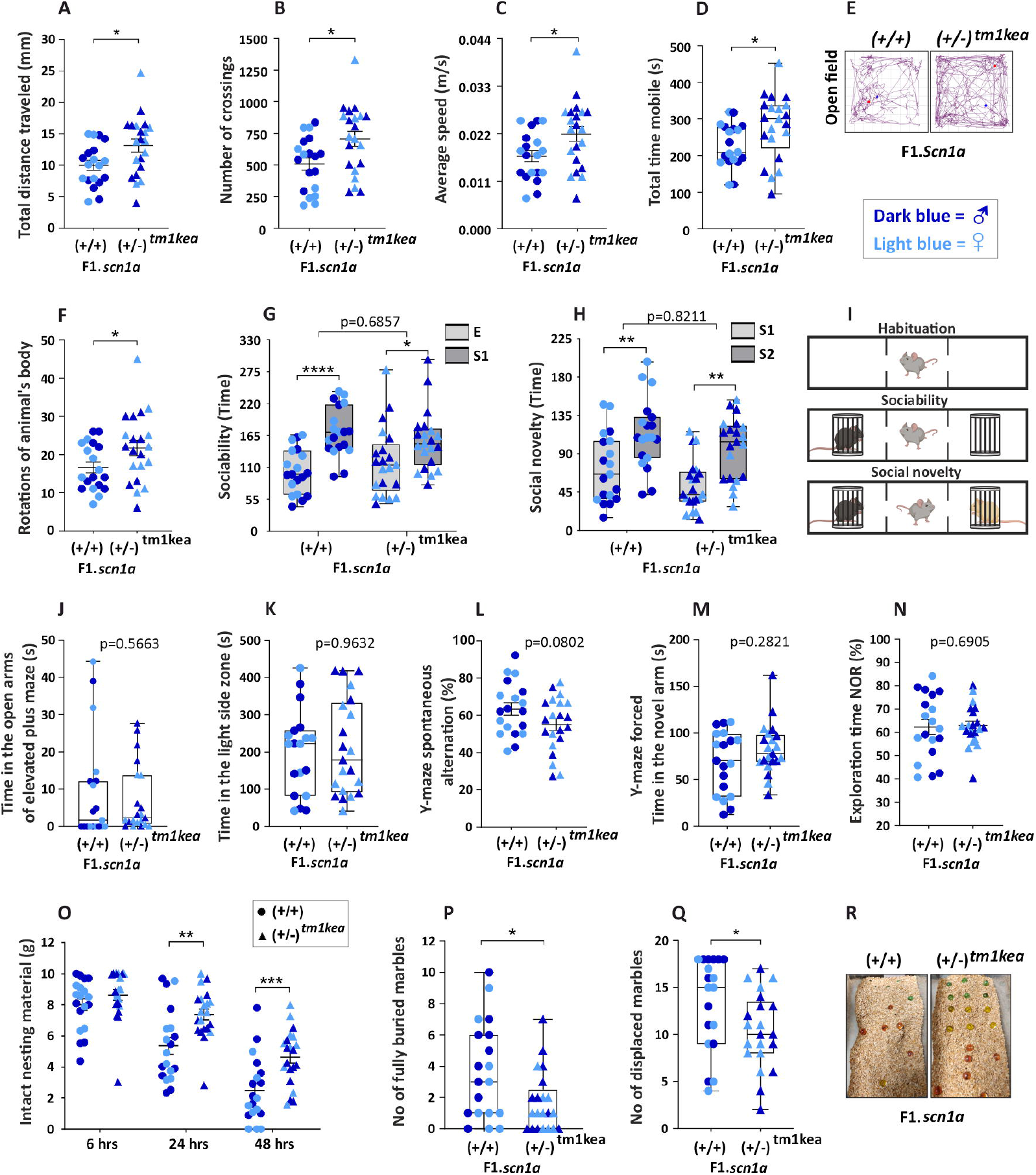
Characterization of long-term neuropsychiatric comorbidities in F1.*Scn1a(+/-)*^*tm1kea*^ mice. Long term comorbidities were investigated across a battery of behavioural assessments with male and female F1.*Scn1a(+/+)*(n=19) or F1.*Scn1a(+/-)*^*tm1kea*^ mice (n = 21/group) around ∼6 months of age. In the open field, F1.*Scn1a(+/-)*^*tm1kea*^ mice A,B,E travelled more, displaying C, higher speed and D, time mobile when compared with F1.*Scn1a(+/+)* mice, indicating hyperactivity. F, F1.*Scn1a(+/-)*^*tm1kea*^ mice displayed a higher number of stereotyped behaviour in the open field indicating autism-like behaviour. G,H,I, However, F1.*Scn1a(+/-)*^*tm1kea*^ mice showed no deficits in sociability or social novelty when compared to F1.*Scn1a(+/+)* mice. Similarly, F1.*Scn1a(+/-)*^*tm1kea*^ mice did not show any anxiety-like behaviour in the J, elevated plus maze and K, Light dark box or memory deficits in L, y maze spontaneous, H, y maze forced or I, in the novel object recognition test when compared to F1.*Scn1a(+/+)* mice. O, F1.*Scn1a(+/-)*^*tm1kea*^ mice displayed poor performance in the nest building assessment when compared to F1.*Scn1a(+/+)*mice indicating poor quality of life. P,Q, F1.*Scn1a(+/-)*^*tm1kea*^ mice also showed a reduced number of fully and displaced marbles when compared F1.*Scn1a(+/+)*mice indicating impaired exploratory behaviour. *p<0.05, **p<0.01, ***p<0.001. A,B,C,D,F,L,N (Student’s t-test), D,J,K,M,P,Q (Mann-Whitney test), G,H (two-way ANOVA), O,(two-way repeated measures ANOVA).

## 4. DISCUSSION

Here we show that F1.*Scn1a(+/-)*^*tm1kea*^ mice display DS-like epilepsy features characterised by sensitivity to febrile seizures at early ages followed by a high occurrence of SRS and SUDEP which diminishes thereafter. SRS and SUDEP were influenced by the time of the day and sex of F1.*Scn1a(+/-)*^*tm1kea*^ mice. Furthermore, F1.*Scn1a(+/-)*^*tm1kea*^ mice displayed some long term comorbidities characterized by hyperactivity, impaired exploratory behaviour but not anxiety, memory or autism-like features and sex differences were not observed. Taken together, these results provide comprehensive assessment of the phenotypes over the lifetime in this model and should facilitate the selection of testing age and sex to improve design and execution of therapeutic and biomarker studies using this DS model.

DS is commonly classified in three different stages accordingly to the clinical manifestations.^7, 8^ The “ febrile stage” of DS is marked by high incidence of hyperthermia sensitivity often resulting in prolonged seizures.^7, 8^ Temperature sensitivity is a conserved feature of mouse models of DS and here we confirmed that F1.*Scn1a(+/-)*^*tm1kea*^ mice present a reduced threshold to the hyperthermia-induced seizures at early developmental stages. These findings are in line with studies showing that F1.*Scn1a(+/-)*^*tm1kea*^ mice display susceptibility to febrile seizures at P18 as also often observed between 6 to 12 months of life in children with DS.^8, 24^ The duration of hyperthermia-induced seizures in F1.*Scn1a(+/-)*^*tm1kea*^ mice has not been previously reported. For the Scn1a+/- knockout DS mouse model, hyperthermia-induced seizures last around 30 s.^25^ Here we found that hyperthermia-induced seizures in P18 F1.*Scn1a(+/-)*^*tm1kea*^ mice fall into two types (short ∼10s and long duration ∼20s) with distinct behaviour which was not influenced by the gender or temperature reached during the hyperthermia challenge. Such a distinct seizure phenotype has not been previously reported in DS mouse models indicating that it is a specific feature only observed in *Scn1a(+/-)*^*tm1kea*^ mouse line.

In children with Dravet, the “ febrile stage” is followed by a “ worsening stage” that extends up to the fifth year of life with increasing SRS frequency and severity. This has also been observed from P21 to P28/P30 in the *Scn1a*+/- knockout mouse model.^8, 23, 25^ Similarly, our results reveal that F1.*Scn1a(+/-)*^*tm1kea*^ mice present a high number of severe SRS over this stage indicating this period features the greatest disturbance of brain excitability. Our data also reveal, however, that female F1.*Scn1a(+/-)*^*tm1kea*^ mice show a higher frequency of SRS than males. Further analyses of sex influence on the severity of SRS also showed that the number of SRS progressing to wild running or jumping (Severity 6) and hindlimb extension (Severity 7) were twice as high in female than in male F1.*Scn1a(+/-)*^*tm1kea*^ mice. Although sex differences in SRS have not been previously reported in DS mouse models and patients, a previous study in F1.*Scn1a(+/-)*^*tm1kea*^ mice reported higher SUDEP rates in female than male DS mice.^26^ Thus, these findings indicate a distinct epilepsy phenotype according to the sex of F1.*Scn1a(+/-)*^*tm1kea*^ mice.

There is increasing evidence that circadian rhythms affect brain excitability.^27^ Previous studies showed that *Scn1a*^R1407X/+^ DS mice experience a peak of SRS ending with death between 18:00 to 19:00 (before the lights went out), higher seizure incidence from 18:00 – 02:00 and during nocturnal period.^28^ Similarly, our findings showed that F1.*Scn1a(+/-)*^*tm1kea*^ mice also experience a peak of SRS before the lights went off (19:00-20:00) and a period of the day with higher seizure incidence (16:00 to 00:00). These results indicate that the time of the day has an influence on the susceptibility to SRS in F1.*Scn1a(+/-)*^*tm1kea*^ mice. It may suggest that network effects of clock-related genes may intersect with the defects arising from *Scn1a* loss. Therefore, pre-clinical studies using F1.*Scn1a(+/-)*^*tm1kea*^ mice should take account a proper monitoring of SRS, particularly if non-continuous monitoring is planned.

Apart from SRS, the worsening stage of DS in humans is marked by high incidence of SUDEP typically preceded by a severe GTCS.^22, 29^ Here, EEG and video monitoring revealed that all deaths experienced by F1.*Scn1a(+/-)*^*tm1kea*^ mice were preceded by a severe GTCS ending with hindlimb extension. SUDEP incidence was highest from P21 to P28, around 80%, and this had no sex bias. The rates of SUDEP are higher than in some studies where video-only monitoring was employed and indicate instrumentation of F1.*Scn1a(+/-)*^*tm1kea*^ mice or daily handling may exacerbate the epilepsy phenotype. Interestingly, neither SUDEP nor changes in background EEG patterns were observed at the beginning of the worsening stage (P21). In contrast, SUDEP observed from P22 to P25 was associated with loss of EEG power during the interictal period in multiple frequency bands including beta, gamma and alpha. Accordingly, a DS mouse model carrying the *Scn1a*^*A1783V*^ missense mutation showed that low EEG power correlates with the risk of premature death during the worsening stage of DS.^30^ Thus, these findings not only characterise the stage 2 of DS in F1.*Scn1a(+/-)*^*tm1kea*^ mice but also reinforce that a severe GTCS and low background EEG power may be an immediate risk factor for SUDEP in DS mice.

The severity and frequency of seizures in humans and mouse models of DS is known to decline in adulthood, often replaced by escalating co-morbidities.^8, 23^ EEG telemetry recordings of F1.*Scn1a(+/-)*^*tm1kea*^ mice confirmed this, with a lower incidence of SRS and SUDEP into adulthood (P36-P49). In naïve F1.*Scn1a(+/-)*^*tm1kea*^ mice followed for 6 months, we found survival rates of ∼60%. More than half of deaths occurred in a short period of time from P20 to P32, tending to decline thereafter until P180 (6 months). These results reinforce previous studies with F1.*Scn1a(+/-)*^*tm1kea*^ mice showing that the critical SUDEP period occurs from P21 to ∼P30, followed by a mitigation period thereafter.^26, 31^ Together, these findings not only characterize the third stage of DS but also reinforces that the epilepsy phenotype in F1.*Scn1a(+/-)*^*tm1kea*^ mice progresses in an age-dependent manner.

A previous study using F1.*Scn1a(+/-)*^*tm1kea*^ mice found that females are more susceptible to SUDEP.^26^ We found a similar increased incidence of SUDEP in naïve female F1.*Scn1a(+/-)*^*tm1kea*^ mice. The most likely explanation is that more frequent seizures are the driver of the higher SUDEP incidence. The mechanism of increased susceptibility to SRS and SUDEP in female F1.*Scn1a(+/-)*^*tm1kea*^ mice is unknown. Factors such as fluctuations in sex hormones or GABA turnover have been found to differ between male and female rodents.^32-36^ Further studies are needed to define the causes of sex differences in DS phenotypes over the different stages of DS in the *Scn1a(+/-)*^*tm1kea*^ mouse model.

DS is also accompanied by multiple long term comorbidities including hyperactivity, social impairments, anxiety and cognitive decline resulting in a poor quality of life.^8, 37^ In the present study, we report the first comprehensive assessment of long-term behavioural phenotypes of F1.*Scn1a(+/-)*^*tm1kea*^ mice during the stabilization stage of DS. Hyperactivity is one of the most consistent findings across different ages of development in individuals with DS.^37^ Accordingly, F1.*Scn1a(+/-)*^*tm1kea*^ mice exhibit hyperactivity when exposed to novel environments with no changes in motor coordination. This is in agreement with the previously reported increase of total distance travelled in the open field in 8 week old male F1.*Scn1a(+/-)*^*tm1kea*^ mice.^26^ Many individuals with DS may also show autistic features including repetitive behaviour and social deficits.^8, 37^ Indeed, we found that F1.*Scn1a(+/-)*^*tm1kea*^ mice exhibited a prominent stereotyped behaviour in the open field arena which may be linked with autism signs in mice. However, F1.*Scn1a(+/-)*^*tm1kea*^ mice did not exhibit social deficits in the three chamber test. These results indicate that the high frequency of repetitive behaviour observed in F1.*Scn1a(+/-)*^*tm1kea*^ mice is probably not an autistic feature, but reflects more likely hyperactive behaviour, as reported in *Scn1a*^WT/A1783V^ mice.^10^

In addition to autistic features, DS patients are reported to have anxiety and especially cognitive deficits.^37, 38^ Three different studies on 6-8 weeks old F1.*Scn1a(+/-)*^*tm1kea*^ mice reported anxiety-like behaviour phenotypes in this DS mouse line.^26, 31, 39^ Similarly, 6 weeks old F1.*Scn1a(+/-)*^*tm1kea*^ mice showed impaired reference and working memory in the radial arm maze.^39^ In contrast, 8 weeks old F1.*Scn1a(+/-)*^*tm1kea*^ mice may display a better novel object recognition memory when compared to WT mice and just slight decline in spatial memory in the Barnes maze test.^39^ Here we consistently found that ∼6 month old F1.*Scn1a(+/-)*^*tm1kea*^ mice did not present any anxiogenic phenotype across the peripheral regions of the open-field, elevated plus maze or light dark box assessment. Furthermore, ∼6 month old F1.*Scn1a(+/-)*^*tm1kea*^ mice did not display spatial reference, working or recognition memory deficits in the forced/spontaneous Y maze or NOR assessment respectively. Together, these results suggest that F1.*Scn1a(+/-)*^*tm1kea*^ mice may experience transient behaviour changes that do not persist into the later stages of life, as similarly reported for the pathophysiology of DS in young adult F1.*Scn1a(+/-)*^*tm1kea*^ mice.^6^

Most DS patients experience a lifelong debilitating condition affecting their daily activities and resulting in a poor quality of life.^40^ In mice, the innate tendency to hide objects in the marble test and nest building activity are relevant measures to compare an individual’s performance in daily life activities.^20^ Indeed, ∼6 month old F1.*Scn1a(+/-)*^*tm1kea*^ mice showed impaired exploratory behaviour as indicated by a reduction in the number of buried and displaced marbles. In addition, F1.*Scn1a(+/-)*^*tm1kea*^ mice displayed a poor performance in the nest building assessment. These results suggest that this epileptic encephalopathy is compromising the welfare and quality of life of F1.*Scn1a(+/-)*^*tm1kea*^ mice as similarly observed in DS patients.

In summary, the present work reports that F1.*Scn1a(+/-)*^*tm1kea*^ mice faithfully reproduce the epilepsy phenotype and select comorbidities of DS, positioning this model as a reliable and versatile tool for the search for new therapies and biomarkers in future pre-clinical tests in DS. However, aspects involving gender and time of the day may influence the epilepsy phenotype of F1.*Scn1a(+/-)*^*tm1kea*^ mice. The findings also point to limitations in the use of older mice in the model to screen for corrective treatments of certain co-morbidities. These factors should be taken into consideration in future studies in *Scn1a(+/-)*^*tm1kea*^ mouse line.

## Supporting information

Characteristics of hyperthermia-induced seizures in F1.Scn1a(+/-)tm1kea mice during febrile stage of DS (P18)

F1.Scn1a(+/-)tm1kea mice do not display anxiety like behaviour in the open field

Sex difference assessment of long-term neuropsychiatric comorbidities in F1.Scn1a(+/-)tm1kea mice

## Authors’ contribution

RRG, CRR and DCH conceived and designed the study and DCH obtained funding. RRG established and performed the management of DS colony with the assistance of JA for genotyping tests and assessment of mice welfare parameters. CRR performed the surgeries for EEG and telemetry recordings. HB performed the EEG background analyses. RRG performed the remaining *in vivo* experiments (hyperthermia challenge, EEG followed by video monitoring and behaviour assessments), data analyses and wrote the initial manuscript. All authors approved the final version of the manuscript.

## ACKNOWLEDGEMENTS

The authors would like to thank Lisa-Ann Byrne and Amaya Sanz-Rodriguez for support with ethical and licensing aspects of the research. This publication has emanated from research supported in part by a research grant from Science Foundation Ireland (SFI) under Grant Number 16/RC/3948 and co-funded under the European Regional Development Fund and by FutureNeuro industry partners. The present study was also supported by funding from the Charitable Infirmary Charitable Trust and F. Hoffman-La Roche Ltd. CRR acknowledges funding from CURE epilepsy.

---

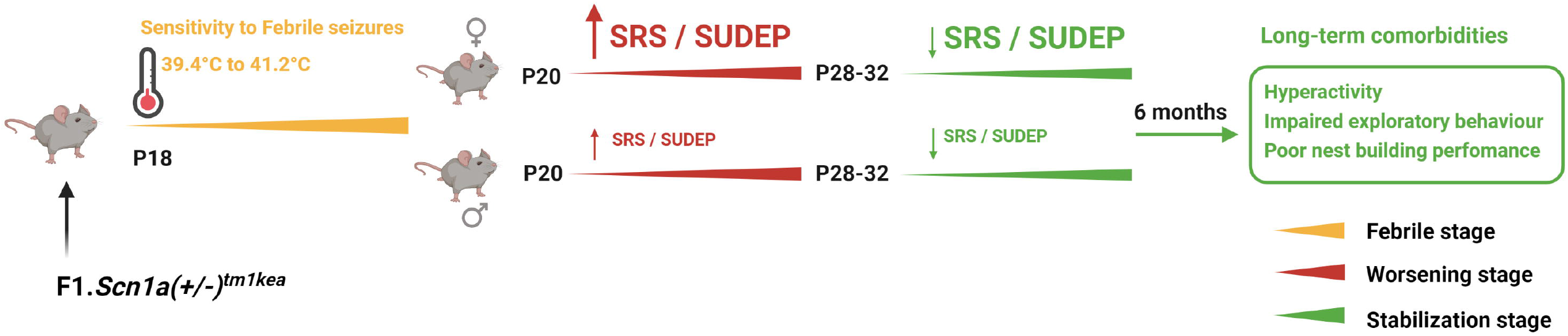

