## Supporting information for "Sexual dimorphism in epilepsy and comorbidities in Dravet syndrome mice carrying a targeted deletion of exon 1 of the *Scn1a* gene"

***Running Title:*** Epilepsy and behaviour phenotype of F1. *Scn1a(+/)*$^{tm1Kea}$ mice

Rogério R. Gerbatin[a,b], Joana Augusto[c], Halima Boutouil[d], Cristina R. Reschke[a,b,e] and David C. Henshall[a,b*]

***Affiliations:***

[a]Department of Physiology and Medical Physics, RCSI University of Medicine and Health Sciences, Dublin, Ireland

[b]FutureNeuro SFI Research Centre, RCSI University of Medicine and Health Sciences, Dublin, Ireland

[c]Department of Physiology, Faculty of Medicine, Trinity College Dublin, Dublin, Ireland

[d]School of Mechanical and Manufacturing Engineering, Dublin City University, Dublin, Ireland

[e]School of Pharmacy and Biomolecular Sciences, RCSI University of Medicine and Health Sciences, Dublin, Ireland

**Corresponding author at:**

### Supporting Information

#### METHODS

##### Genotyping of F1.*Scn1a(+/-)*$^{\mathrm{tm1Kea}}$ mice

Tissue was collected by tail snipping before P7. DNA was extracted and subject to PCR using KAPA2G Fast Genotyping Mix (KAPA Biosystems, UK). Primers used for genotyping are described in the table below.

##### Hyperthermia-induced seizures

Hyperthermia-induced seizure threshold assay was performed at P18 as previously described.[1] First, the mouse was gently hand-restrained in a supine position with tail lifted. Then, a temperature probe (RET-4, physitemp) covered with Vaseline was inserted into the rectum and taped on the tail, to keep it in place throughout the procedure. Later, animals were placed into a Plexiglas box with an infrared heat lamp (HL-1, physitemp, Clifton, New Jersey) positioned above and the rectal probe attached to a TCAT-2DF thermocontroller (physitemp, Clifton, New Jersey). Mice were held at 37.5°C for 5 min to become accustomed to the chamber and then core body temperature was gradually elevated by 0.5°C every 2 min until a seizure (GTCS) occurred or 42.5 °C was reached. Body temperature was then elevated by 0.5 °C every 2 min until a seizure occurred or 42.5 °C was reached. If the mouse had a seizure, the heating process was stopped immediately to cooled down the mouse to 37°C in a cold metal surface. Animals not presenting seizures were held for 3 min until 42.5°C and then the heat lamp was turned off. Mice remained 5 minutes into the chamber for observation of any eventual late occurrence of seizure before they are removed, cooled down and considered seizure free. Seizure severity was classified according to the Racine scale scoring system with few modifications.[2, 3]

#### Acute video-EEG recordings of SRS and death

At P21, another cohort of mice were injected via i.p with buprenorphine (0.3mg/ml) and placed in an adapted stereotaxic frame under anaesthesia (isoflurane/oxygen 5% for induction and 3% for maintenance). Body temperature was maintained by a feedback-controlled heat blanket. After topical application of EMLA cream 5%, a midline scalp incision was performed, and three screw electrodes were implanted and secured with dental cement and special glue. The screw electrodes were placed bilaterally to the midline over the cerebral cortex followed by the reference electrode positioned over the nasal sinus. After surgery, animals were immediately placed in an incubator at 33°C and monitored for 30 min. Once fully recovered, mice single housed were connected to the lead socket of a swivel commutator, which was connected to a brain monitor amplifier for EEG digital recordings. Gel diet was added in the cage and video-electroencephalography (vEEG) recordings were performed from 12:30 p.m to 18:30 p.m (6 hours/day) followed by video monitoring from 18:30 p.m to 12:30 p.m (18 hours/day) until P28.

#### EEG background analyses

EEG background analysis was performed using MATLAB. Three hours of EEG was selected from the interictal period from each file and the extracted data was organized following the mice gender (Male and Female) and the disease (WT/Het). The signal analysis was done using wavelet transform and FFT (Fast Fourier Transform).

#### Video-EEG monitoring of SRS and death

Immediately after acute vEEG recordings, single housed mice in their home cages were transferred to a room equipped with a high resolution, infrared video cameras (Hikvision). Continuous digital videos were recorded at 30 fps and stored in a Dell PC workstation. Video's recording was offline review at 16x speed using VSplayer (version 6.0.0.4) software and suspected seizures were reviewed at real time. Duration of seizures were defined from the beginning to end of behavioural convulsion. Seizure severity was classified according to the Racine scale scoring system with few modifications as previously mentioned in the section 2.3.1 (Hyperthermia induced-seizures).[2, 3] When a mouse was found dead in a cage, the video was reviewed to determine time of death and whether it was preceded by a severe GTCS ending with full hindlimb extension.

#### Long term continuous video-EEG recordings of SRS, SUDEP and survivals analyses in naïve mice

At P36, another group of mice were implanted with telemetry devices for long term vEEG recordings. The telemetry device (Model: F20-EET, Data Systems International) was inserted subcutaneously along the dorsal flank. Four electrodes were connected to skull-mounted screws, 2 references electrodes were placed in the nasal sinus, one electrode was placed over the right cerebral cortex and another electrode placed over the left cerebral cortex. Continuous vEEG monitoring (telemetry system; Data Sciences International, USA) was performed 24/7 for 2 weeks until P49. Finally, another batch of naïve mice was used to monitor the survival rates from P0 to 6 months of age, period that long-term comorbidities assessment started as described below.

#### Behavioral experiments

Animals reaching 5 to 6 months of age were submitted to a battery of behavioral tests designed with multiple resting days interval and according to increasing order of invasiveness. 2 weeks before starting the behavioral assessment, animals were moved to the animal unit equipped with all behavioral apparatus to get familiarized with new environment. 1 week before starting the battery of behavioral tests, all mice were handled during 5 min in three different days with interval of 24 hours in the behaviour unit to get familiarized with the researcher and environment. After each behaviour session, apparatus and objects used were cleaned with 70% ethanol to avoid any potential bias. All behavioral sessions were video recorded by an overhead Logitech Webcam C270 (720p) and analysed with ANY-maze video tracking system (Version 6.32) when appropriated.

#### Forced alternation test (Y-maze)

Spatial reference memory was assessed as previously described.[4] The mouse was placed into the end of the start arm, facing the wall and away from the centre. In the trial, mouse was allowed to explore during 5 min the two arms of the Y-maze, while entry into the third arm was blocked. After the sample trial, the mouse was returned to its home cage for a 60 min inter-trial interval. In the trial 2, the block in arm 3 was removed and the mouse was again placed into the start arm, and then allowed to access all three arms of the maze. If a mouse climbed on the maze wall, it was immediately returned into the abandoned maze arm. An arm entry was recorded when 85% of a mouse's body entered the arm. Time in Novel Arm [%] was defined as the time spent in the novel arm divided by the time spent in all arms during the first minute of the retrieval trial T2. Forced Alternation [%] was defined as the percent of mice entering first the novel arm during T2. Mice with less than three arm entries in the first minute of T2 were excluded from the analysis.

**Novel object recognition**

Recognition memory was measured using the novel object recognition task as previously described.[5, 6] During habituation (Day 1), mice were placed on the middle of the open empty arena (27,3 × 27,3 × 27,3 cm) to freely explore it for 10 min. Twenty-four hours later (Training phase – Day 2), two identical objects in opposite quadrants of the arena were adhered to the floor. Mice were placed in the arena for 10 min and later returned to their home cage. After 60 min (short-term memory; STM) or 24h (long-term memory), two dissimilar objects were presented (a familiar and a novel one) and time spent exploring each object was measured during 10 min. Two falcon® tubes (50 mL, 30 × 115 mm), filled with clean bedding and a lego toy and a Corning® U-shaped polystyrene cell culture flasks (75 cm2) filled with clean bedding were used as a objects. To avoid any potential bias, the locations of objects for each session were counterbalanced according to mice genotypes. A recognition index calculated for each animal in STM task was expressed by the ratio TB/(TA + TB) [TA = time spent exploring the familiar object A; TB = time spent exploring the novel object B] and in LTM task was expressed by the ratio TC/(TA + TC) [TA = time spent exploring the familiar object A;TC = time spent exploring the novel object C]. To avoid confounds by lingering olfactory stimuli and preferences, the objects and the arena were cleaned with 70% ethanol after each animal was tested. Video recordings were manually offline reviewed by an experienced researcher. Exploration was defined as sniffing or touching the objects with the nose and/or forepaws. Sitting on or turning around the objects was not considered exploratory behaviours. Data acquired during the habituation stage of NOR in the open field was used quantify the spontaneous locomotion and anxiety-like behaviour.

**Spontaneous alternation test (Y-maze)**

Spatial working memory test was conducted using a symmetrical Y-maze (35 cm x 5 cm x 60 cm) with the floor and walls made of white and clear Plexiglas respectively as previously described.[4] For this test, the Y-maze was rotated by 45° and the distal cues (geometrical marks) were differed between the forced alternation tests. During a single 5 min trial, mice freely explored all three arms of the Y-maze containing different geometrical marks (black and/or white) on the wall of each arm. The start arm was varied across genotypes to avoid placement bias. Mice that climbed on the maze walls were immediately returned to the abandoned arm. Spontaneous alternation [%] was defined as consecutive entries in 3 different arms (ABC), divided by the number of possible alternations (total arm entries minus 2). Mice with less than 8 arm entries during the 5-min trial were excluded from the analysis.

**Three chamber tests of social preference**

The three-chamber apparatus (Ugo Basile Sociability Cage/code 46503) is a transparent plexiglass box (60 x 40 x 22 (h) cm) divided into three identical chambers separated by two transparent partitions. Each partition has a square opening (5 x 5 cm) in the bottom centre that offer free access to any partitions. Three chambered social preference test consists in 3 sessions respectively named as habituation, sociability and social novelty session performed as previously described with modifications.[7] Test mice were confined in the centre chamber at the beginning of each session. In the habituation session, a test mouse was placed in the centre of the three-chamber unit to freely explore the area during 10 min. Then, for the sociability session, a stranger age- and gender-matched C57BL/6J mouse (S1) that had never been exposed to the test mouse was placed in one of the two wire cups leaving the opposite wire cup empty. The location of the S1 alternated between the two side chambers across test mice. During the social novelty phase, a stranger mouse (S2) was enclosed in the wire cup that had been empty during the sociability phase. Thus, the test mouse would have the choice between a mouse that was already familiar (S1) and a new stranger mouse (S2). Exploration of an enclosed mouse or a wire cup was defined as when a test mouse oriented toward the wire cup with the distance between the nose and wire cup less than 1 cm. C57BL/6J mouse were habituated in the three chamber apparatus 1 week before the tests during 10 minutes into the wire cup in three different days to avoid any potential stress that could interfere in the test day.

**Elevated plus maze (EPM)**

The elevated plus maze apparatus comprised two open arms (25 cm length x 5 cm width) and two closed arms (25 cm length x 5 cm width x 16 cm height walls to enclose the arms) placed in the centre of the testing room, 50 cm above the floor. The test consisted of a single trial lasting 5 minutes. Mice were individually placed in the centre facing one of the open arms. The number of entries into the open and closed arms and the time spent was recorded. These parameters were used to calculate the index of anxiety-like behaviour.

**Marble burying**

Marble burying was used to assess the innate exploratory behaviour of mice.[8] The test was conducted in opaque Perspex cages (35 cm × 16,5 cm × 14 cm) containing normal bedding (5 cm deep) overlaid with 18 glass marbles (15 mm diameter) equidistant in a 3 × 6 arrangement. For testing, a single mouse was placed in the cage with the lid closed for 30 min. At the end of the trial, the mouse was removed from the cage, and a researcher blinded to genotype counted the number of marbles buried. Marbles with more than 2/3 of its surface covered was counted as buried. Bedding was replaced, and marbles were cleaned with 70% ethanol, rinsed with water, and dried between trials.

#### Light / Dark box

The light/dark task was performed as previously described [9]. The light-dark box consists of an arena (0.5m x 0.5m), divided equally into a safe, dark compartment and an illuminated aversive compartment. The dark compartment is enclosed on all sides with a small opening to allow the animal to travel into the light compartment. Mice are placed individually in the centre of the light compartment facing the opening to the dark compartment. Parameters evaluated are: (a) cumulative time spent in light (aversive) and dark (non-aversive) compartments is recorded during a 10 min session; (b) number of entries into each compartment as defined by all four paws being placed in that compartment; (c) number of head entries (nose-pokes) into the light compartment; (d) distance travelled by the animal in each compartment. An increase or decrease in total time spent in the light compartment reflects, respectively, a decrease or increase in 'anxiety', with distance travelled reflecting level of 'activity', and number of head entries reflecting a 'decision-making' variable.

#### Rotarod

Motor coordination and balance were evaluated on rotarod (Harvard apparatus model LE 8500), accelerating from 4 to 40 rpm in 300 s [10]. The mouse ability to maintain balance and keep pace with a rotating road were evaluate during 5 min over three trials with interval of 15 min. Latency to fall was recorded for each trial.

#### Nest building

Single-housed mice were transferred into a new cage with 100g of soft cob bedding containing 10g of nest-building material cut into square (Whatman filter paper, grade 201, Sigma).

After 6, 24, and 48 h, the weight of intact nest material was assessed to determine the nest building ability as previously described [11].

#### Tables

**Table S1.** Primers used for genotyping of F1.*Scn1a(+/-)*$^{tm1Kea}$ mice.

| Primer | Catalogue no. | Sequence (5'→ 3') |
|---|---|---|
| Common | 19078 | AGT CTG TAC CAG GCA GAA CTT G |
| Wildtype reverse | 19181 | CCC TGA GAT GTG GGT GAA TAG |
| Mutant reverse | oIMR2088 | AGA CTG CCT TGG GAA AAG CG |

#### Figure Legends

**Figure S1.** Characteristics of hyperthermia-induced seizures in F1.*Scn1a(+/-)*$^{tm1kea}$ mice during febrile stage of DS (P18). A, Sex of F1.*Scn1a(+/-)*$^{tm1kea}$ mice does not influence on the threshold to trigger a seizure during the hyperthermia challenge. B, Duration of hyperthermia-induced seizures in F1.*Scn1a(+/-)*$^{tm1kea}$ mice does not correlate with temperature threshold which seizure occurs. A, Permutation test, B Spearman rank-order correlation.

**Figure S2.** F1.*Scn1a(+/-)*$^{tm1kea}$ mice do not display anxiety like behaviour in the open field. Male and female F1. *Scn1a(+/+)*(n=19) or F1.*Scn1a(+/-)*$^{tm1kea}$ mice (n = 21/group) around ~6 months of age were submitted to the open field test. A,B, F1.*Scn1a(+/-)*$^{tm1kea}$ mice displayed higher number of entries in the centre and corner zone of open field without C,D, increasing the time spend in these areas when compared to F1. *Scn1a(+/+)* mice, indicating only hyperactivity traits but not anxiety. E,F,G,H Subgroup analysis of male (n=11) and female (n=10) F1.*Scn1a(+/-)*$^{tm1kea}$ mice revealed no significant difference in any anxiety-like behaviour parameters. Mann-Whitney test *p<0.05.

**Figure S3.** Sex difference assessment of long-term neuropsychiatric comorbidities in F1.*Scn1a(+/-)*$^{tm1kea}$ mice. Long term comorbidities was investigated across a battery of behaviour assessment between male (n=11) and female (n=10) F1.*Scn1a(+/-)*$^{tm1kea}$ mice around ~6 months of age. Analysis of male and female F1.*Scn1a(+/-)*$^{tm1kea}$ mice revealed no significant difference in any behaviour parameters. A,B,C,D, Male and female F1.*Scn1a(+/-)*$^{tm1kea}$ mice displayed no difference in parameters related to the spontaneous locomotion in the Open field and E,F,G autism-like behaviour in the open field and three chamber test. Similarly, no significant difference was found between male and female F1.*Scn1a(+/-)*$^{tm1kea}$ mice in parameters related to anxiety in the H, elevated plus maze, I, Light dark box or memory deficits in J, Y maze spontaneous, K, Y maze forced and L, in the novel object recognition. Similar performance between male and female F1.*Scn1a(+/-)*$^{tm1kea}$ mice was observed in the M, nest building assessment and N,O, exploratory behaviour in the marble burying test. A,B,C,,E,J,K,L (Student's t-test), F,G (Two-way ANOVA), D,H,I,N,O (Mann-Whitney test) and M (Two-way repeated measures ANOVA). *p<0.05

**Movie S1.** Short hyperthermia-induced seizure experienced by F1.*Scn1a(+/-)*$^{tm1kea}$ mice.

**Movie S2.** Long hyperthermia-induced seizure experienced by F1.*Scn1a(+/-)*$^{tm1kea}$ mice.

**Movie S3.** Typical severe SRS experienced by F1.*Scn1a(+/-)*$^{tm1kea}$ mice.
