## Supplementary figures and images for "Sexual dimorphism in epilepsy and comorbidities in Dravet syndrome mice carrying a targeted deletion of exon 1 of the *Scn1a* gene"

### Characteristics of hyperthermia-induced seizures in F1.Scn1a(+/-)tm1kea mice during febrile stage of DS (P18)

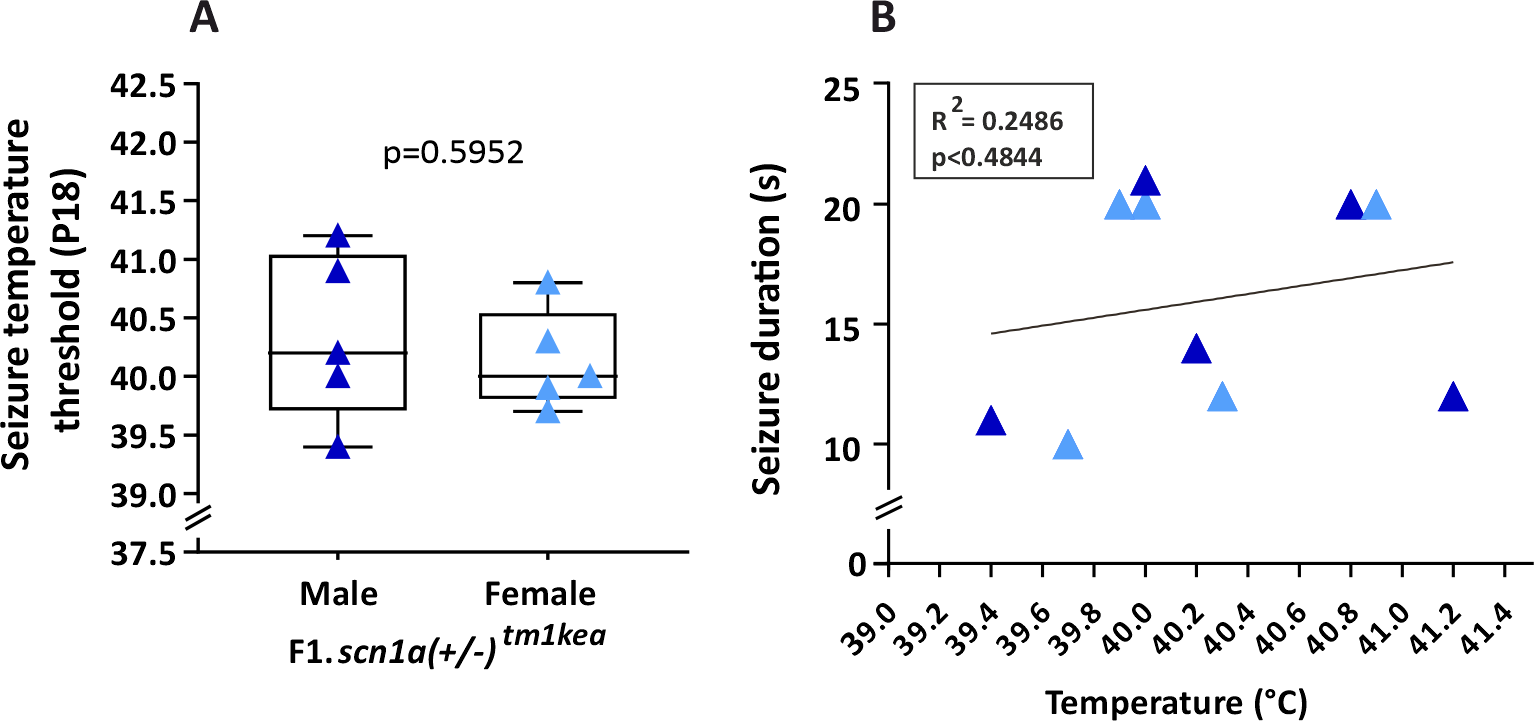

### F1.Scn1a(+/-)tm1kea mice do not display anxiety like behaviour in the open field

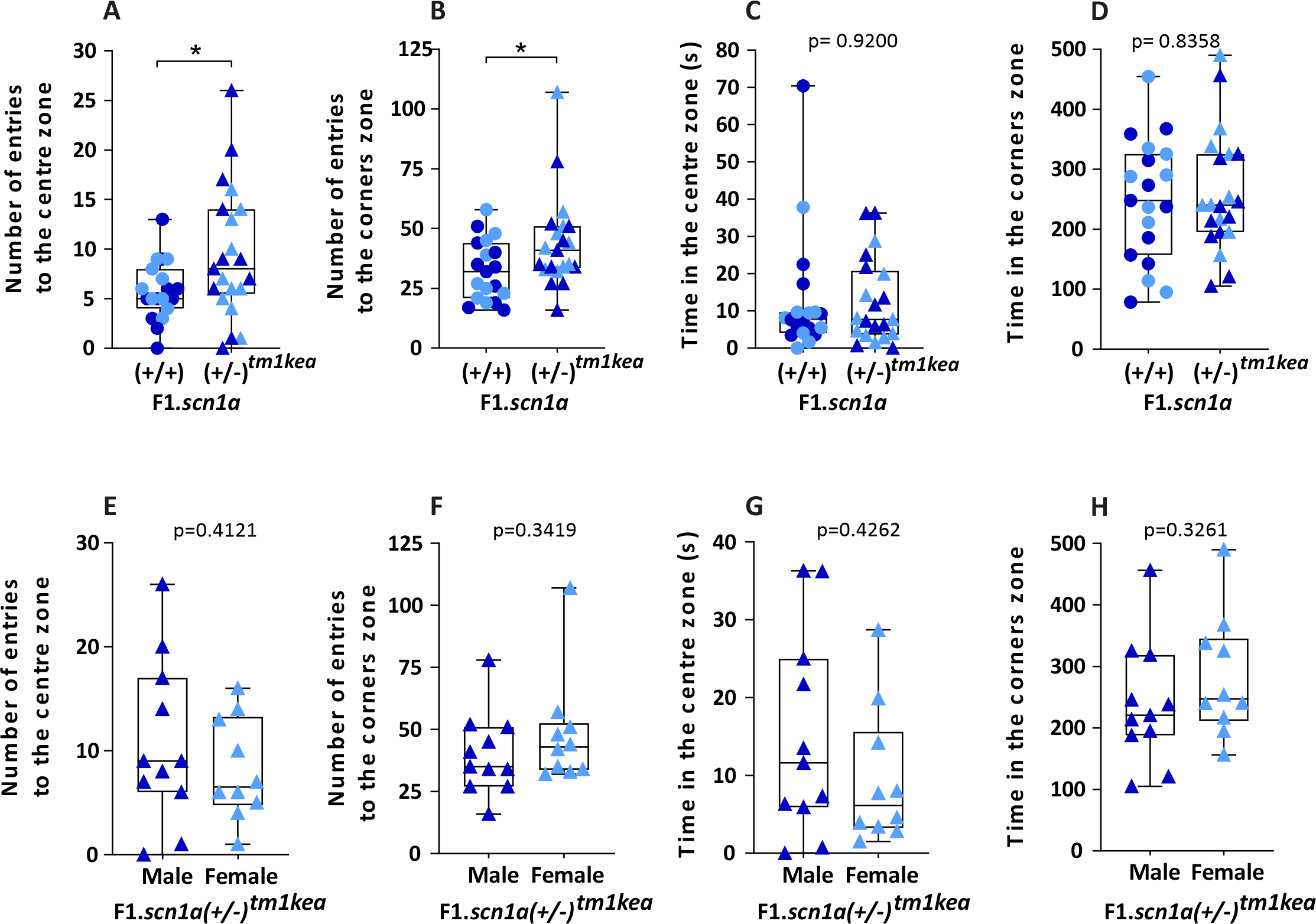

### Sex difference assessment of long-term neuropsychiatric comorbidities in F1.Scn1a(+/-)tm1kea mice

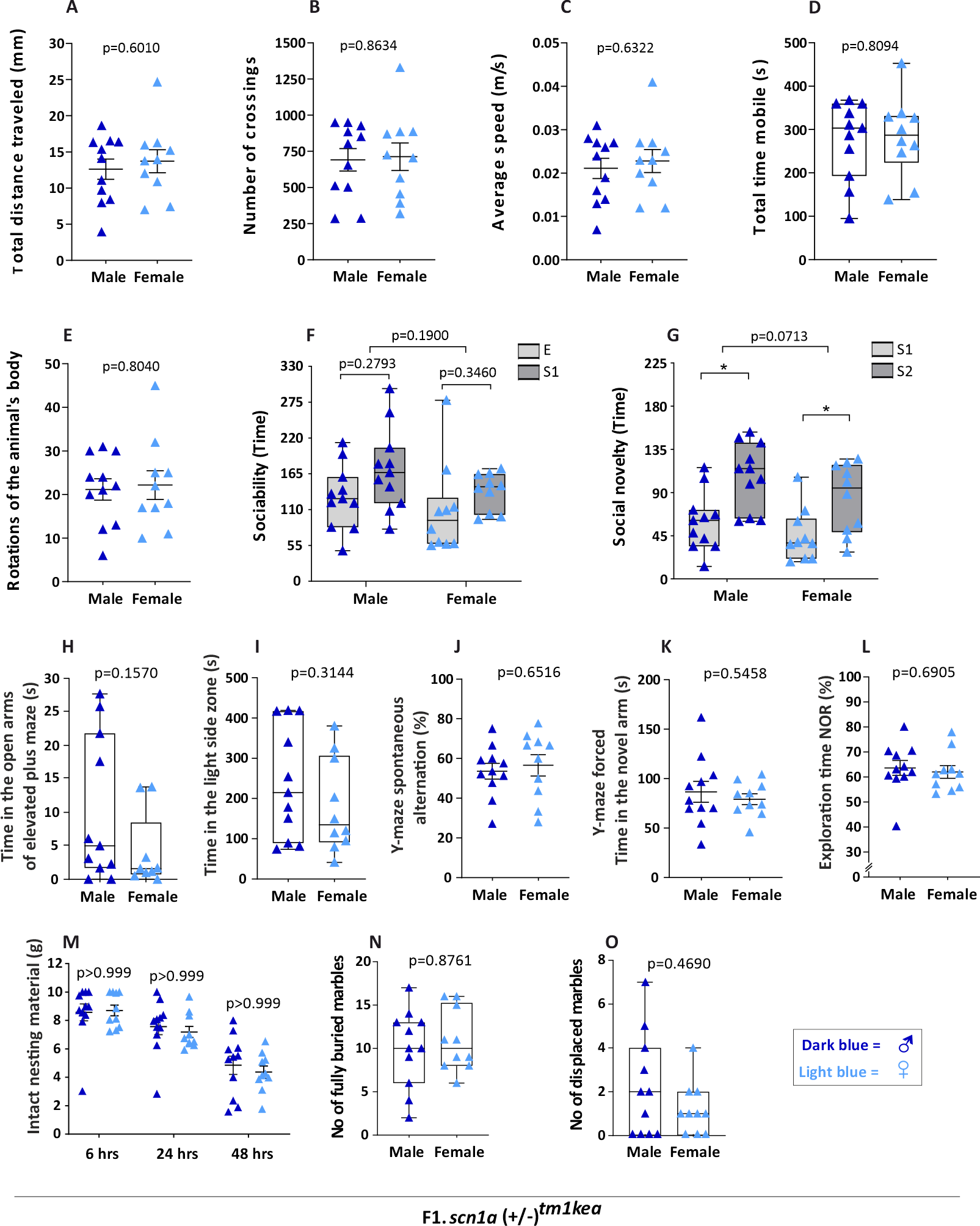
